# DNA-metabarcoding uncovers the diversity of soil-inhabiting fungi in the tropical island of Puerto Rico

**DOI:** 10.1101/025668

**Authors:** Hector Urbina, Douglas G Scofield, Matias Cafaro, Anna Rosling

**Affiliations:** Department of Evolutionary Biology, Uppsala University, Norbyvägen 18D, Uppsala 752 36, Sweden; Uppsala Multidisciplinary Center for Advanced Computational Science, Department of Information Technology, Uppsala University, Box 137, Uppsala 751 05, Sweden; Department of Biology, University of Puerto Rico, Mayagüez 00681-9000, Puerto Rico

**Keywords:** Communities, Dikarya, high-throughput sequencing, internal transcribed spacer, molecular operational taxonomic unit

## Abstract

Soil fungal communities in tropical regions remain poorly understood. In order to increase the knowledge of diversity of soil-inhabiting fungi, we extracted total DNA from top-organic soil collected in seven localities dominated by four major ecosystems in the tropical island of Puerto Rico. In order to comprehensively characterize the fungal community, we PCR-amplified the internal transcribed spacer 2 (ITS2) fungal barcode using newly designed degenerated primers and varying annealing temperatures to minimize primer bias. Sequencing results, obtained using Ion Torrent technology, comprised a total of 566,613 sequences after quality filtering. These sequences were clustered into 4,140 molecular operational taxonomic units (MOTUs) after removing low frequency sequences and rarefaction to account for differences in read depth between samples. Our results demonstrate that soil fungal communities in Puerto Rico are structured by ecosystem. Ascomycota, followed by Basidiomycota, dominates the diversity of fungi in soil. Amongst Ascomycota, the recently described soil-inhabiting class Archaeorhizomycetes was present in all localities, and taxa in Archaeorhizomycetes were among the most commonly observed MOTUs. The Basidiomycota community was dominated by soil decomposers and ectomycorrhizal fungi with a distribution strongly affected by local variation to a greater degree than Ascomycota.

## 1. Introduction

Recent studies using next generation sequencing (NGS) technologies have increased our understanding of the diversity of soil-inhabiting fungi enormously. Most surveys were carried out in the Northern Hemisphere, more specifically in Europe and North America (e.g., Clemmensen et al. 2013; Lentendu et al. 2014; Menkis et al. 2015), however a few investigations targeting soil-inhabiting fungi in tropical regions (e.g., Kemler et al. 2013; Tedersoo et al. 2014). Initial surveys indicate that tropical biomes contain the greatest species richness of soil fungi (McGuire et al. 2013), though across different biomes, soil fungal diversity appears to be similarly structured by abiotic and biotic factors (McGuire et al. 2013; Tedersoo et al. 2014). We do not yet have more extensive sampling from a single tropical region, and more efforts are needed to extend the characterization of tropical soil fungal communities.

The Caribbean tropical island of Puerto Rico exhibits a rich diversity of flora and fauna in just 8,900 km^2^ (Liogier and Martorell 2000; Gannon et al. 2005). This landmass together with Cuba and Hispaniola form the Greater Antilles of the West Indies, which are all fragments of old continental crust (Ricklefs and Bermingham 2008). Puerto Rico has a complex geographical history due to several periods of separation and rejoining with the other Greater Antilles (Ricklefs and Bermingham 2008). Six major ecosystems have been identified: littoral zone forest, semi-deciduous subtropical dry forest, tropical-and subtropical-moist forest, and subtropical rain forest (Helmer et al. 2002).

Previous studies in Puerto Rico have aimed to characterize fungi adapted to hypersaline environments using classic molecular and culture-based approaches (Cantrell et al. 2007; Cantrell and Duval-Perez 2012) and basidiomycete ectomycorrhizal fungi (Miller et al. 2000). More recently, the diversity of soil fungi has been characterized using NGS techniques at three localities in El Yunque National Forest (NF) (Tedersoo et al. 2014). Consequently, soil-inhabiting fungal communities remain largely uncharacterized throughout this island. The aim of this work is to provide a greater insight into the diversity of soil-inhabiting fungi across the major ecosystems in Puerto Rico.

## 2. Materials and methods

### 2.1. Soil samples and total DNA extraction

Three plots were sampled at each of seven localities representing four major ecosystems in Puerto Rico (Montane and Subtropical Wet Forest, Subtropical Dry Forest and Littoral) (Fig. 1, Table 1). The top-organic layer of plant debris and rocks was removed in an approximately 1 m^2^ grid and a table spoon of soil was scooped from each corner and the center of an internal 0.50 m^2^ grid. In the field, soils were pooled in a sterile plastic bag and manually homogenized for 1 min. Two sub-samples of approximately 0.5 g were added into separate 2.0 ml microtube containing 750 μl of lysis buffer (Xpedition^TM^ Soil/Fecal DNA miniprep, Zymo Research Corporation, Irvine, California, USA), followed by cell disruption for 30 s using a TerraLyser^TM^ (Zymo Research Corporation), carried as well in the field. Microtubes were stored at 4 °C until the following steps of the DNA extraction was carried out in the laboratory within one month of sampling, following the manufacturer’s protocol. DNA concentration and integrity was verified in 0.8 % agarose gel electrophoresis on 0.5 % Tris Acetate-EDTA buffer (Sigma-Aldrich, St. Louis, Missouri, USA) stained with 1 × GelRed^TM^ (Biotium Inc., Hayward, California, USA). DNA from a pure culture of *Neurospora crassa* was included as a positive control.

**Fig. 1 –.**
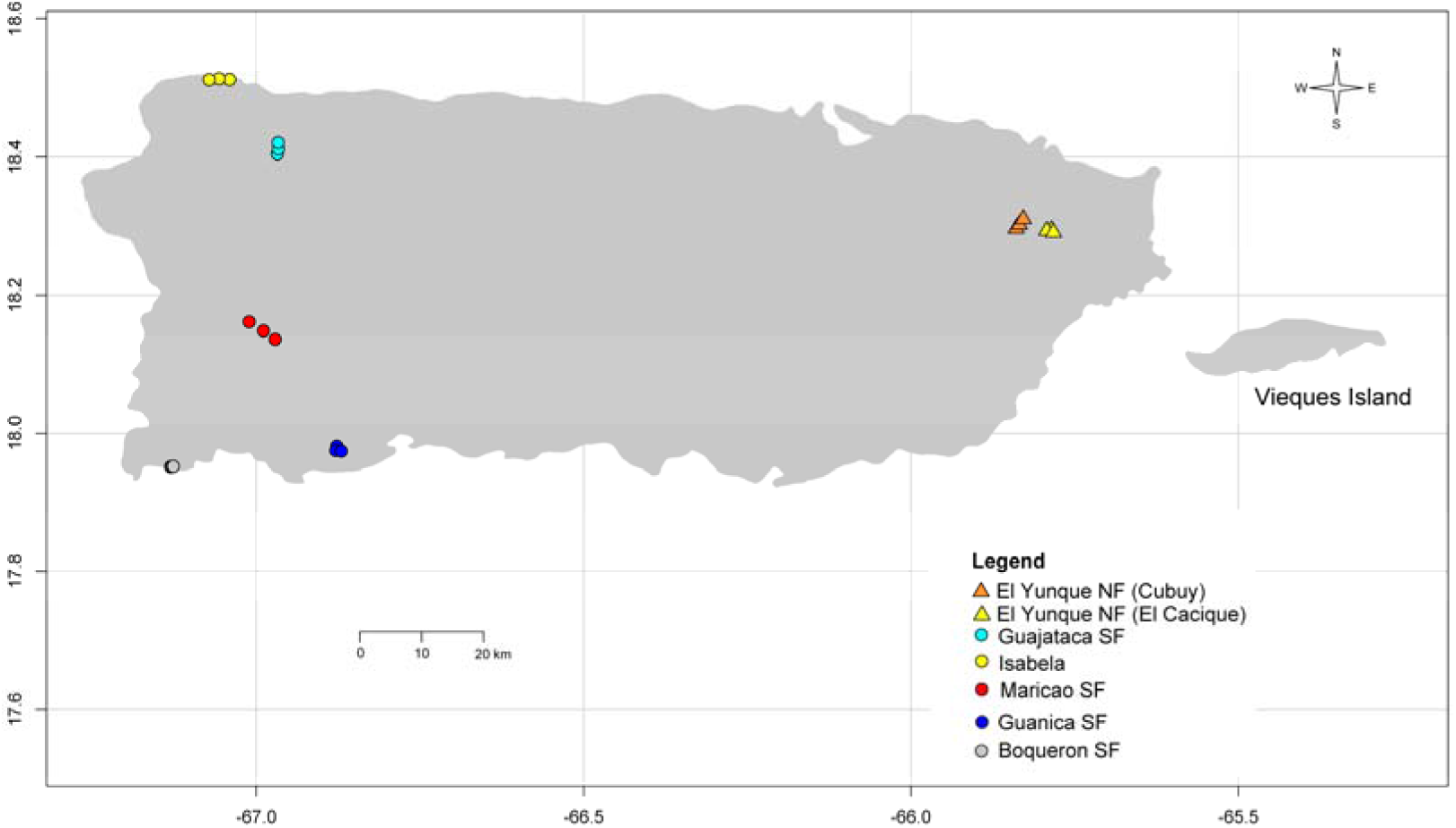
Map of Puerto Rico showing localities of sampling sites. National forest (NF) in triangle; and states forest (SF) in circle.

**Table 1 –.**
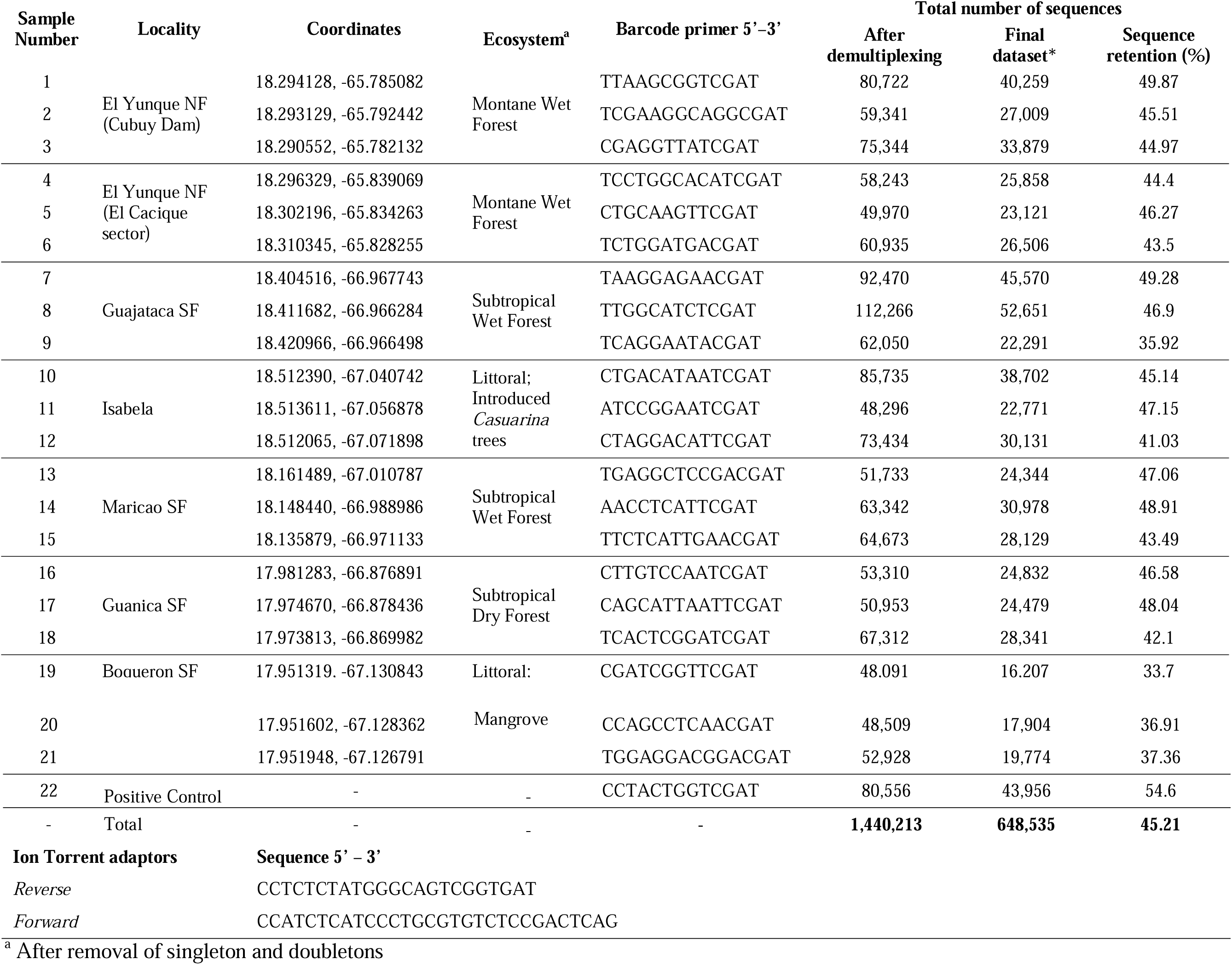
Collecting localities in Puerto Rico, barcode primers used and sequencing results after demultiplexing and removal of singleton and doubletons.

### 2.2. Polymerase Chain Reaction (PCR) and Ion Torrent library preparation

A fragment of the 5.8S, the internal transcribed spacer 2 (ITS2) and a fragment of the large subunit (LSU) of the rRNA genes was amplified using primers gITS7 *forward* (Ihrmark et al. 2012) and modified ITS4m *reverse* (5’-TCCTC[**C/G**][**G/C**]CTTATTGATATGC-3’), with both primers containing adequate barcode sequences for single-ended amplification (Table 1). The ITS locus is broadly accepted as a taxonomic barcode for *Fungi* (Schoch et al. 2012) and among ITS1 and ITS2 there are no significant differences in their power of discriminating species between fungal groups (Blaalid et al. 2013). Modifications on the reverse primer ITS4 (White et al. 1990) were included to reduce its known bias against the soil-inhabiting fungal class Archaeorhizomycetes (Schadt and Rosling 2015).

The PCR mixes were comprised of 10-20 ng of soil DNA, 1 × SSoAdvanced^TM^ Universal SYBR^®^ Green Supermix (Bio-Rad Laboratories, Hercules, California, USA), and 0.8 nM of each primer in a final volume of 20 μl. PCR amplifications were carried out in a CFR96 Touch^TM^ Real/Time PCR Detection system (Bio-Rad Laboratories) following the protocol, 10 min pre-denaturation at 95 °C, 1 min DNA denaturation at 95 °C, 45 s at three independent annealing temperatures (50, 54 and 58 °C) to reduce primer bias (Schmidt et al. 2013), 50 s of extension at 72 °C and 3 min final extension at 72 °C. We used a quantitative PCR (qPRC) to adjust the number of cycles for each plate between 23-27, in order to ensure that the reactions were maintained within linear amplification. To reduce the chance of altering the relative abundance of fungi by biasing against long reads, we used tag primers directly thereby avoiding an extra nested-PCR run. All reactions were carried out in duplicates, and all six runs were combined before purification using the ZR-96 DNA Clean & Concentrator™-5 (Zymo Research Corporation). DNA concentration was quantified on duplicates using the Quant-iT™ PicoGreen^®^ dsDNA Assay Kit (Life Technologies Corporation, Carlsbad, California, USA) on a TECAN F500 microplate reader and the DNA integrity was checked by electrophoresis in 2 % agarose gel 0.5 × TAE buffer. A sequencing library was prepared by pooling 35 ng DNA from each sample, loaded onto a 318 chip for PGM Ion Torrent sequencing technology (Life Technologies Corporation) and sequenced in the facilities at Uppsala Genome Center (Uppsala University, Sweden).

### 2.3. Assembly and taxonomy of molecular operational taxonomic units (MOTUs)

We obtained a demultiplexed dataset comprising 1,440,213 sequences, with 1,359,657 sequences corresponding to samples and 80,556 to the positive control (Torrent Suite v 4.0.4) (Table 1, Supplementary Fig.S1A). The software mothur v1.33.3 (Schloss et al. 2009) was used for sequence trimming of the FastQ files (*Q* ≥ 25 in sequence average, minimum length 150 bp) (Table 1, Supplementary Fig. S1B). To improve the accuracy of clustering and assigned taxonomy of the MOTUs, we used only the variable ITS2 locus, by trimming the fragments of 5.8S and LSU loci using the software ITSx v1.0.9 (Bengtsson-Palme et al. 2013). After this step, all sequences shorter than 80 bp and non-fungal sequences were removed; consequently our dataset was reduced to 648,535 sequences of fungal ITS2 locus exclusively (Supplementary Fig. 1C).

The ITS2 fungal sequence dataset was *de novo* clustered using the software TSC (Jiang et al. 2012) under the following parameters: data type 454 (Ion Torrent and 454 LifeSciences sequencing technology possesses similar sequencing errors (Yang et al. 2013)), single linkage clustering algorithm, recommended for ITS locus in fungi (Lindahl et al. 2013), sequence identity 0.97 (suitable cutoff for species delimitation in fungi (Blaalid et al. 2013)). We tested three cutoff values for high (HB) and low abundance (LB) sequences (100, 500 and 1000) and obtained 18,961, 18,293 and 18,292 MOTUs, respectively for each cutoff. Because cutoffs 500 and 1000 converged to similar total number of MOTUs, we choose 500 for further analysis. Singletons and doubletons were then discarded. The final ITS2 dataset contained 608,300 sequences and 4,378 MOTUs, representing 97.3 % of sequences and 23.9 % of MOTUs from the ITS2 dataset, a reduction that allowed for robust analyses (Supplementary Fig. 1D). Ion Torrent sequencing technology can prematurely truncate sequences (Salipante et al. 2014), thus the longest sequences representing each MOTUs were selected using the script *pick_rep_set.py* in QIIME v1.8.0 (Caporaso et al. 2010). The initial taxonomic assignment of MOTUs was performed using BLASTn 2.2.29+ (Madden 2002) with parameters E-value = 1e^-10^ and word size = 7, against the reference database 2015-03-02 UNITE + INDS (Koljalg et al. 2013) with only trimmed ITS2 region as described above. Unassigned MOTUs were blasted against the nucleotide database at NCBI and sequences of the ITS2 hits were extracted along with their respectively taxonomy using custom scripts deposited at https://github.com/douglasgscofield/add-to-Qiime-DB. For any sequences with taxonomic levels not matching those in the 2015-03-02 UNITE + INDS database, we corrected their taxonomy using Index Fungorum (http://www.indexfungorum.org). We also added Archaeorhizomycetes representative sequences (Menkis et al. 2014).

The final taxonomic identification of each representative sequence was corrected based on 97, 90, 85, 80, 75 and 70 % of sequence identity for assigning MOTUs with names of a species, genus, family, order, class, or phylum, respectively, as proposed by Tedersoo et al. (2014).

The positive control using DNA of *N. crassa* was used to verify the entire protocol from sampling and tag primer arrangement to clustering and taxonomic assignment. The majority of sequences (95 %) in this sample were clustered into a unique MOTU that was identified as *N. crassa*. The error of 5 % was distributed across 53 MOTUs. Eleven of the MOTUs were unique of the positive control (288 sequences) the rest represents MOTUs also identified in the main dataset and showed variable sequence abundance from 1-243 sequences. We concluded that this error originates predominantly from the demultiplexing step and it did not affect the abundance of any particular MOTU significantly (less than 0.1%). The positive control was eliminated from further analysis. The script *compute_core_microbiome.py* in Qiime was used to account for the difference in sequence depth among samples by the rarefaction method using the lowest number of sequences (11,455) found in one of the samples as recommended by (McMurdie and Holmes 2014). This resulted in a reduction to 4,140 MOTUs. Sequence representatives for each MOTU that satisfied GenBank requirements are available under the accession numbers (KT241043-KT245134). Sequence data and abundance information per soil sample and locality are available in the Supplementary online File S1.

### 2.4. Statistical analyses

Sequences abundance plots were generated using the library *phyloseq* (McMurdie and Holmes 2013) in R v3.0.2 (http://www.r-project.org). In order to address differences in MOTUs composition and diversity, rarefaction curves, Chao-1 and Shannon diversity indexes and the non-metric multidimensional scaling (NMDS) based on Bray Curtis distance were computed in the R package *ampvis* (http://madsalbertsen.github.io/ampvis/). To address the statistically significance of the variables studied here the multivariate analysis of variance *Adonis* implemented in Qiime was used. Double clustering analysis based on the 100 most abundant MOTUs was carried out using the R script *heatmap.2* (http://www.insider.org/packages/cran/gplots/docs/heatmap.2). The g-Test of independence was used to calculate the significance of association of MOTUs with locality and ecosystem and corrected by Bonferroni and False Discovery Ration test with 1000 permutations, all as implemented in the Qiime script *group_significance.py*.

## 3. Results

We identified a total of 4,140 MOTUs at different taxonomic levels after rarefaction: 559 to species (13.5 %), 965 to genus (23.3 %), 919 to family (22.2 %), 785 to order (19.0 %), 354 to class (8.6 %), 548 to phylum (13.2 %) and only 10 MOTUs (0.2 %) remained unclassified but assigned to kingdom fungi.

In terms of taxonomic abundance, Dikarya dominates the community of soil-inhabiting fungi in Puerto Rico (Fig. 2A, Supplementary online file S1). MOTUs classified in Ascomycota were the most abundant (2,967 MOTUs; 79.6 % sequences) followed by Basidiomycota (1,022 MOTUs; 17.5 % sequences). Much less abundant fungi included Glomeromycota (206 MOTUs; 1.6 % sequences;), Chytridiomycota (35 MOTUs; 0.2 % sequences), Zygomycota (68 MOTUs; 0.2 % sequences), and Cryptomycota (67 MOTUs; 0.3 % sequences) and 0.6 % of the sequences remained unidentified.

**Fig. 2 –.**
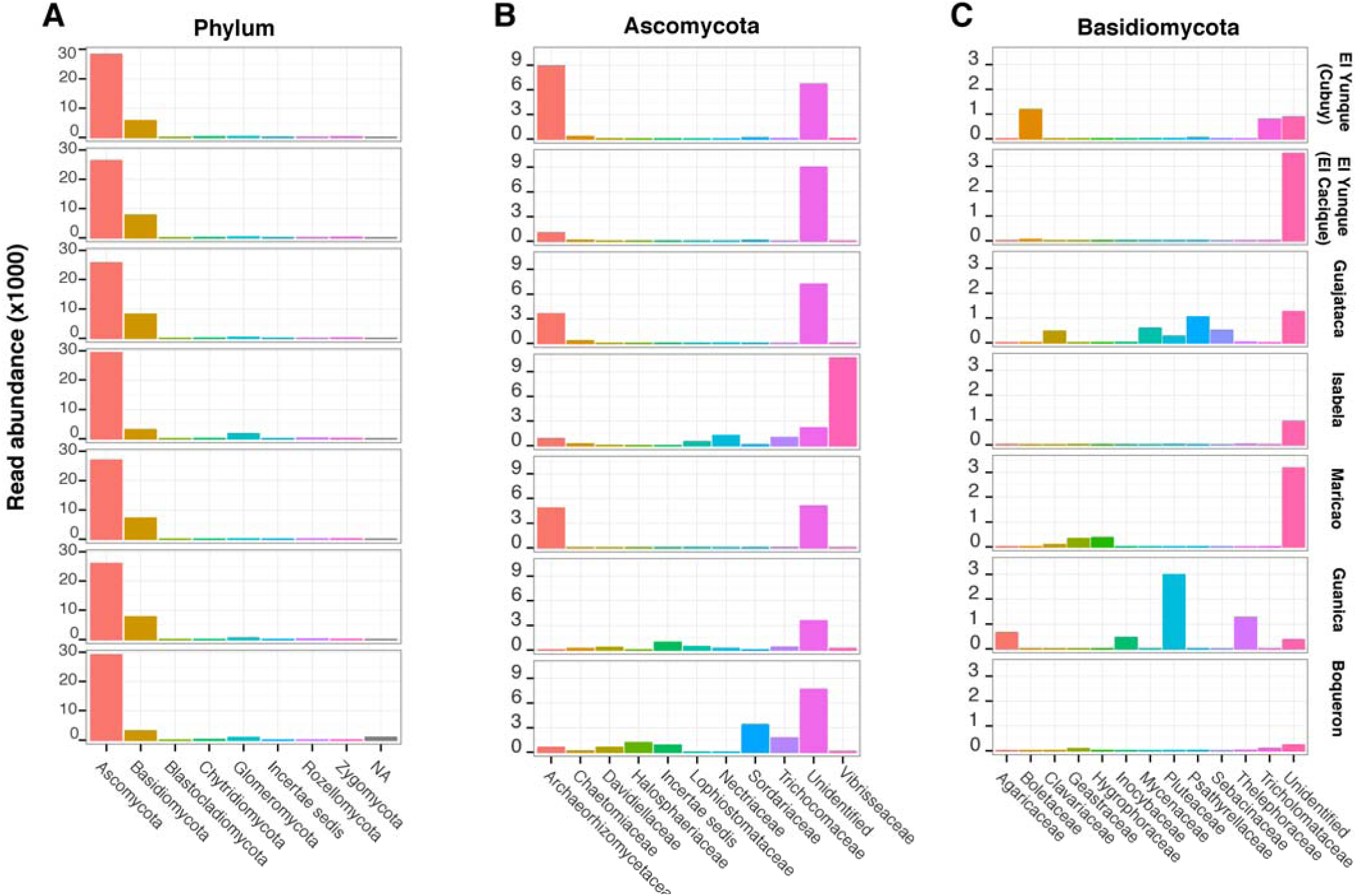
Read abundance of MOTUs at each locality: A) Phyla; B) Families in Ascomycota; C) Families in Basidiomycota.

Among Ascomycota the classes with higher sequence abundance belong to Archaeorhizomycetes (20.9 %), Sordariomycetes (13.9 %), Eurotiomycetes (11.4 %), Dothideomycetes (10.6 %) and Leotiomycetes (10.6 %) (Figs. 2B, 3). Among ascomycetes, 26 MOTUs were found across all localities: *Archaeorhizomyces* (4 MOTUs), *Pestalotiopsis* (2 MOTUs), *Aspergillus, Bionectria, Chaetomium, Cladosporium, Cylindrocladium, Haematonectria, Lasiodiplodia, Neurospora, Penicillium, Phialocephala* and *Talaromyces* (1 MOTU each) (Supplementary online File S1). Focusing on the most abundant class Archaeorhizomycetes, a total number of 50,379 sequences were assigned to the class, these grouped into 190 MOTUs (4.6 % of the total MOTUs) (Supplementary online File S1).

**Fig. 3 –.**
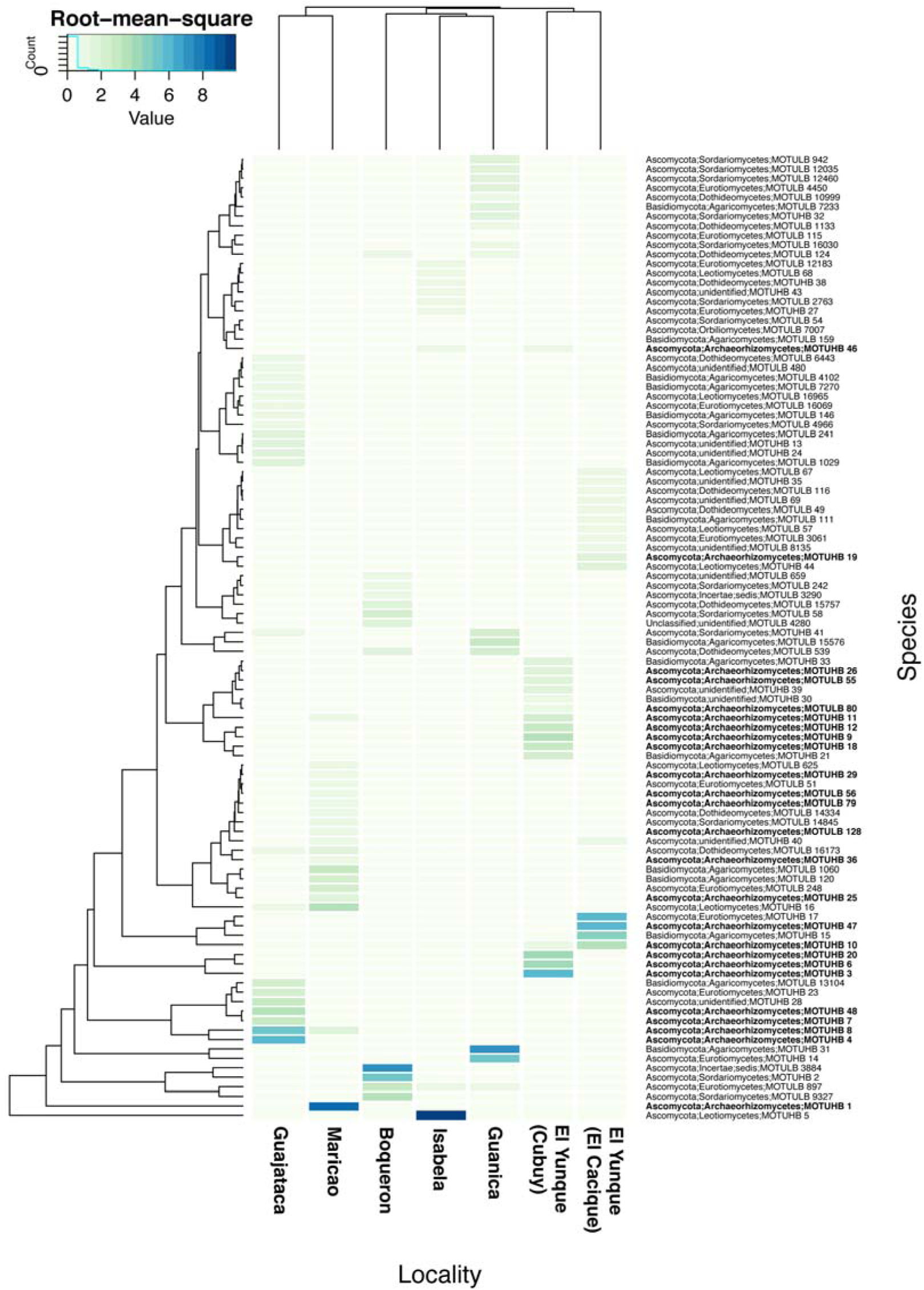
Double complete clustering based on Euclidean distance of the 100 most abundant of MOTUs. Top tree: Locality. Left tree: MOTU. MOTU abundance. Archaeorhizomycete MOTUs are indicated in bold type.

Among Basidiomycota, the class Agaricomycetes was the most diverse with 669 MOTUs (15.3 % sequences) followed by unclassified Basidiomycota 207 MOTUs (1.3 %, Fig. 2C). The most diverse families were Agaricaceae, Clavariaceae, Mycenaceae, Thelephoraceae and Psathyrellaceae, most members of which are saprophytic (Supplementary online File S1). Ectomycorrhizal fungi (EMF) classified in the Boletaceae, Inocybaceae, Pluteaceae and Sebacinaceae (108 MOTUs) (Supplementary online File S1) were also detected in Puerto Rico (Fig. 2C). Guajataca was the locality in which basidiomycetes were most abundant (134 MOTUs; 8,103 sequences) followed by Maricao (256 MOTUs; 7,700 sequences). The most abundant MOTU was classified as unidentified Pluteaceae (MOTUHB 31; 3,326 sequences). It was detected at two localities, most abundantly in Guanica (3,044 sequences) with markedly fewer sequences in Guajataca (255 sequences). This pattern of strongly site-specific abundances was observed among the most abundant basidiomycete MOTUs: 15, 1060, 120, 21, 13104, 33 and others (Fig. 3, Supplementary online File S1). The majority of basidiomycete MOTUs had fewer than 20 sequences in total (746 MOTUs, 73.1 %) and no Basidiomycota MOTU were found at all localities, in contrast to Ascomycota.

Among all sites, Glomeromycota was most abundant in Isabela (46 MOTUs; 1,281 sequences), the littoral *Casuarina* site. In contrast to nearly all other localities, its abundance was consistent among samples (Fig. 2A).

The MOTU accumulation curves indicate high variation in number of MOTUs per sample (200-800 MOTUs in total, Supplementary Fig. S2) and localities (900-1200 MOTUs in total, Fig. 4). The observed MOTU accumulation curves continued to accumulate total alpha diversity, while both the Chao-1 richness and Shannon diversity measures reached or approached a plateau in the majority of localities (Fig. 4) as well as samples (Supplementary Fig. S2). This indicates that we have a significant portion of soil fungal diversity at our samples, while rare taxa likely remains undetected. Among localities the number and proportion of common MOTUs are shown in the Table 2. On average, the number of MOTUs shared between pairs of localities was 20.8 %. Over all the percentage of shared MOTUs was higher between localities from the same ecosystem than between localities from different ecosystem.

**Fig. 4 –.**
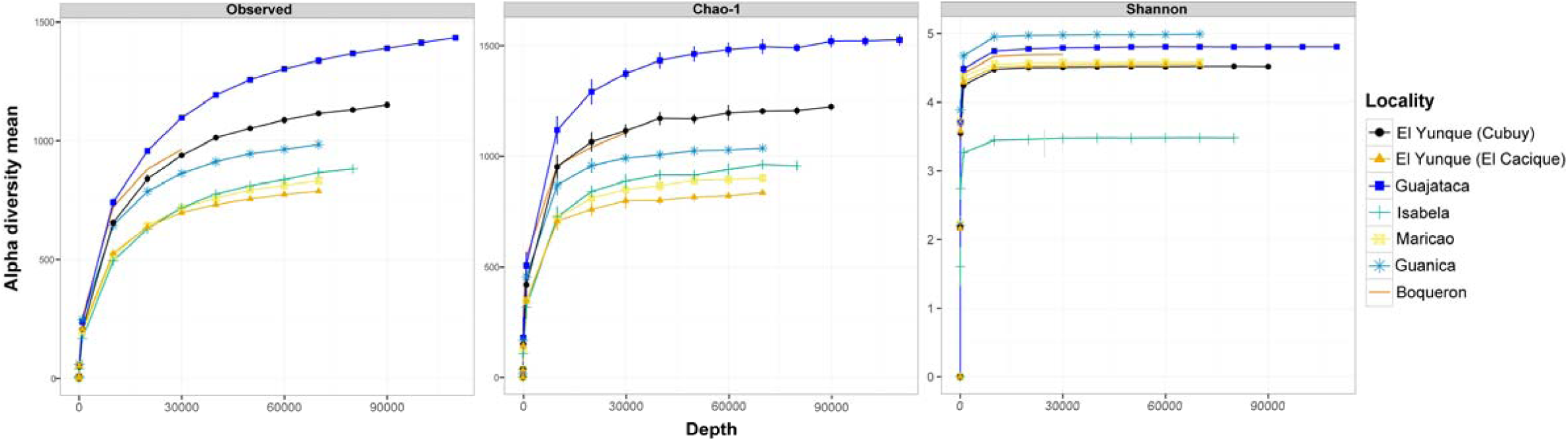
MOTU accumulation curve of alpha diversity by locality among observed species and the Chao-1 and Shannon diversity indices.

**Table 2 –.**
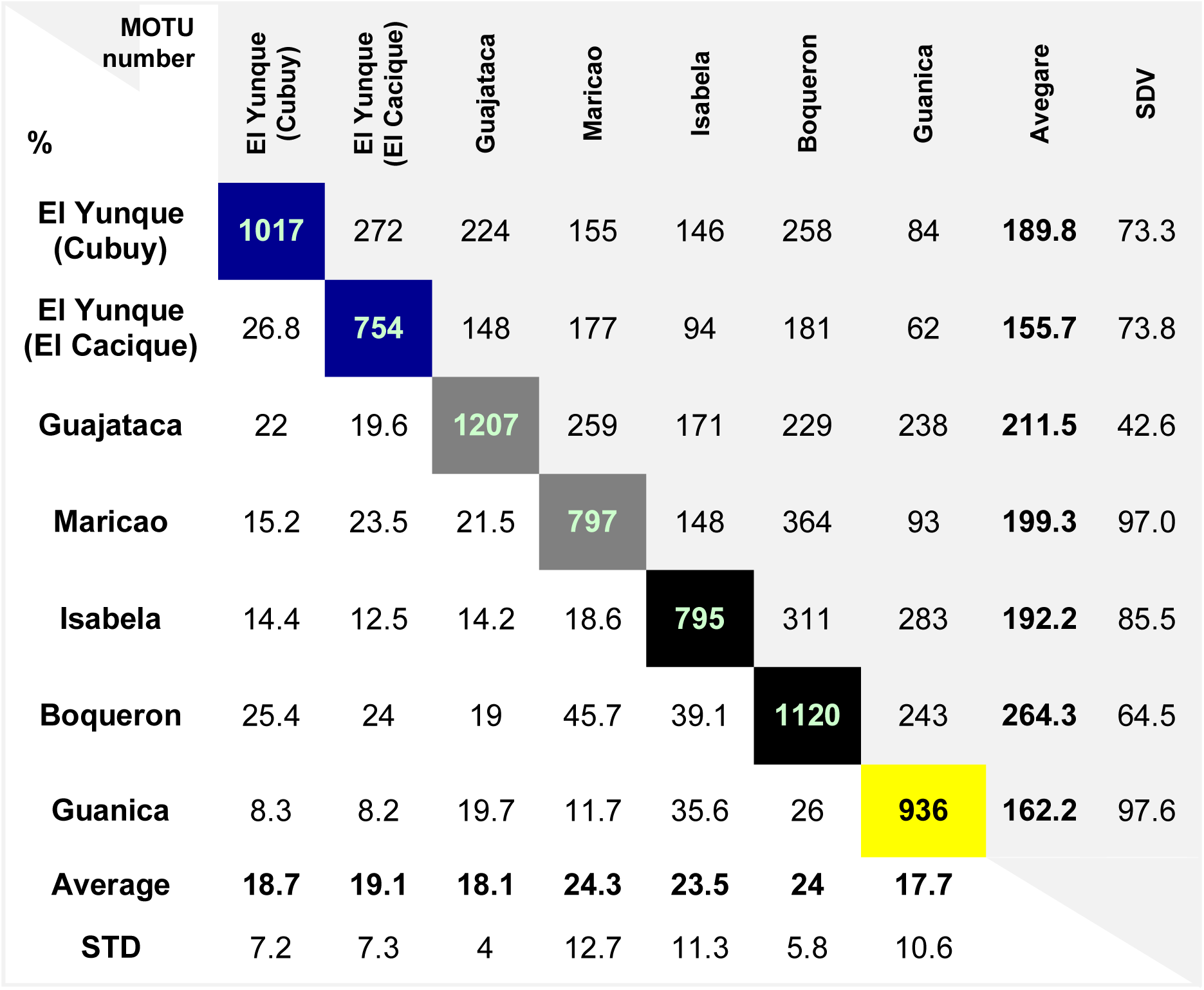
Total number and percentage of common MOTUs per locality. Cell background by ecosystem: Montate Wet (dark blue)-, Subtropical Moist (dark grey), Littoral (black), and Subtropical Dry (yellow). Standard deviation (SDV)

| MOTU<br>number | El Yunque<br>(Cubuy) | El Yunque<br>(El Cacique) | Guajataca | Maricao | Isabela | Boqueron | Guanica | Avegare | SDV |
| --- | --- | --- | --- | --- | --- | --- | --- | --- | --- |
| % |  |  |  |  |  |  |  |  |  |
| El Yunque<br>(Cubuy) | 1017 | 272 | 224 | 155 | 146 | 258 | 84 | 189.8 | 73.3 |
| El Yunque<br>(El Cacique) | 26.8 | 754 | 148 | 177 | 94 | 181 | 62 | 155.7 | 73.8 |
| Guajataca | 22 | 19.6 | 1207 | 259 | 171 | 229 | 238 | 211.5 | 42.6 |
| Maricao | 15.2 | 23.5 | 21.5 | 797 | 148 | 364 | 93 | 199.3 | 97.0 |
| Isabela | 14.4 | 12.5 | 14.2 | 18.6 | 795 | 311 | 283 | 192.2 | 85.5 |
| Boqueron | 25.4 | 24 | 19 | 45.7 | 39.1 | 1120 | 243 | 264.3 | 64.5 |
| Guanica | 8.3 | 8.2 | 19.7 | 11.7 | 35.6 | 26 | 936 | 162.2 | 97.6 |
| Average | 18.7 | 19.1 | 18.1 | 24.3 | 23.5 | 24 | 17.7 |  |  |
| STD | 7.2 | 7.3 | 4 | 12.7 | 11.3 | 5.8 | 10.6 |  |  |

We used non-metric multidimensional scaling (NMDS) of the 21 samples to illustrate how fungal community composition differed with locality and ecosystem (Fig. 5). This analysis showed that the soil fungal communities are distinct depending on both the locality (*r^2^* = 0.844, *P* < 0.01) and ecosystem (*r^2^* = 0.781, *P* < 0.01) (Fig. 5), reflecting the MOTU-specific analysis and confirmed using clustering analysis (Fig. 3). This same pattern was also recovered analyzing the Ascomycota and Basidiomycota communities separately (Ascomycota, locality: *r^2^* = 0.831, *P* < 0.01; ecosystem: *r^2^* = 0.7794, *P* < 0.01), Basidiomycota, locality: *r^2^* = 0.796, *P* < 0.01; ecosystem: *r^2^* = 0.532, *P* < 0.02; Supplementary Fig. S3).

**Fig. 5 –.**
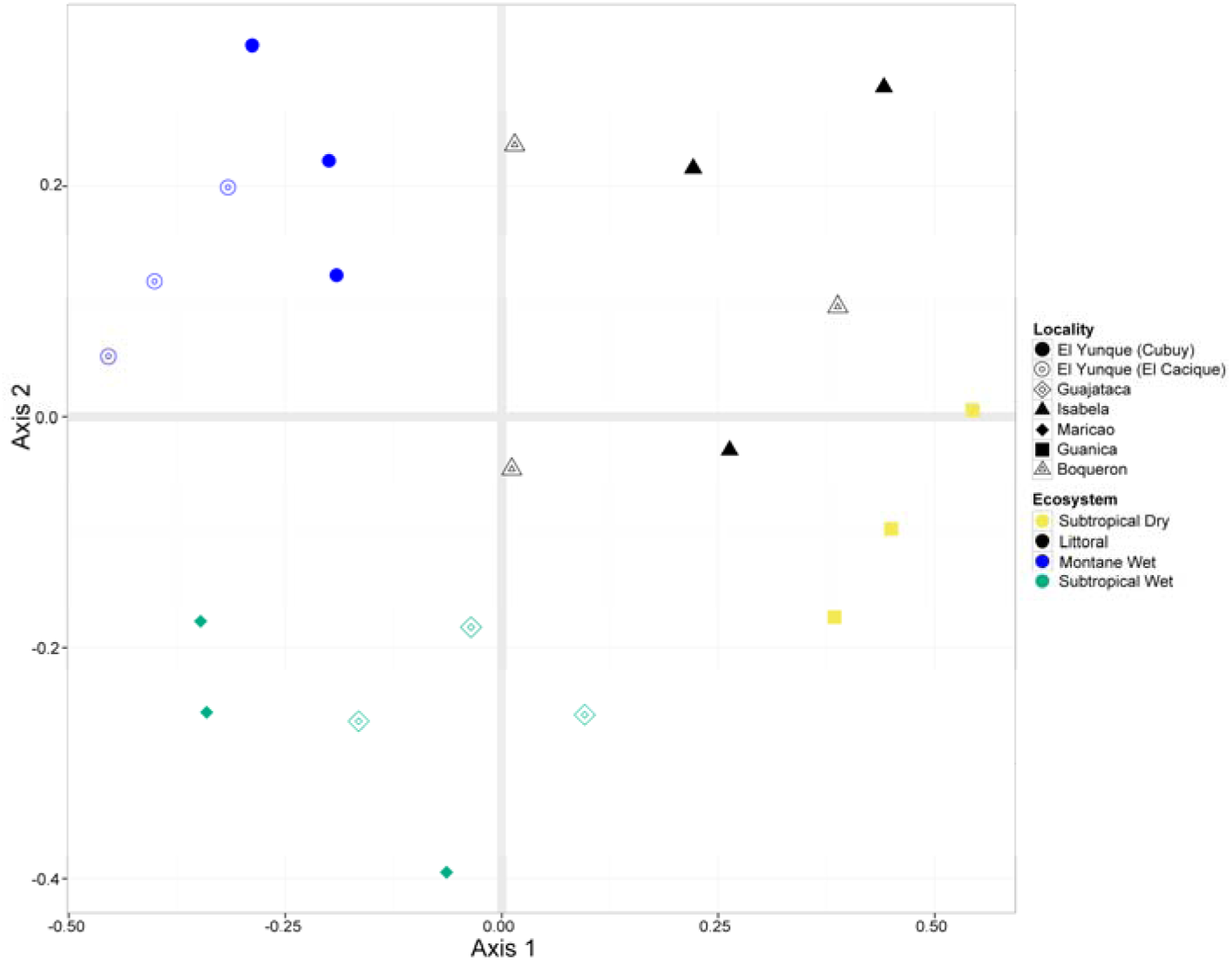
Non-metric multidimensional scaling (NMDS) of all MOTU abundances in each sample, categorized by locality and major ecosystem.

## 4. Discussion

### 4.1. Fungal identification based on MOTUs

Most of the MOTUs were identified to family or higher taxonomic levels (63.2 %). The lack of ITS2 reference sequences of fungi from tropical regions as well as possible fungal endemism in Puerto Rico are likely reasons for the identification of a high number of MOTUs resolved only to high taxonomic levels such as phylum or class. This phenomenon is common in many recent studies, e.g., in Menkis et al. (2015) 13 of the 30 most common MOTUs were identified only to phylum, and in Toju et al. (2014) 10 of the 25 most common MOTUs were identified only to phylum.

We found that the majority of MOTUs are unique to each sampling locality (Table 2). Specifically, we reported 1,499 MOTUs from El Yunque (Cubuy and El Cacique) while independently Tedersoo et al. (2014) found 1,652 fungal MOTUs in the same locality but at different sampling sites (Supplementary Fig. S4). Between the two studies, 264 MOTUs (17.8 % of our MOTUs) were found in both based on a blast sequence similarity search using 97% similarity against our MOTU dataset. Even though Tedersoo et al. (2014) and our study are independent and used different sampling and DNA extraction protocols, both captured similar MOTU richness in the same ecosystem despite high local variation and relatively low MOTU overlap. In the future, field surveys should take into account the degree of spatial separation between sites (McGuire et al. 2013) to increase the amount of expected overlapping species between individual samples in order to capture a more complete set of fungal species.

### 4.2. Soil-inhabiting Ascomycota in Puerto Rico

We found that Ascomycota dominated soil fungal communities; this is in accordance with earlier studies demonstrating that this phylum dominates soil fungal communities in the majority of ecosystems worldwide (Toju et al. 2014; Menkis et al. 2015). Soil fungi in this phylum include decomposers of organic matter and many taxa establish symbiotic relationships with plant roots (Smith and Read 2008). In contrast to our results Tedersoo and co workers (2014) found greater abundance of Basidiomycota relative to Ascomycota in tropical soils, including samples from El Yunke N.P. in Puerto Rico. Such discrepancies demonstrate that while NGS has greatly increased our knowledge about soil fungal communities much work remains until these are properly described even at high taxonomic rank.

Class Archaeorhizomycetes was the most species rich ascomycetous group found in Puerto Rico. This is in agreement with previous observations that the richness of Archaeorhizomycetes is highest in tropical regions (Tedersoo et al. 2014). As a result of our use of un-biased primers we observed higher sequences abundance and hence detected more species in the class compared to Tedersoo et al (2014). Only eight (4.4%) of the MOTUs assigned to Archaeorhizomycetes were identified as species previously detected among environmental sequences. This is because available sequences assigned to the class have been obtained predominantly from studies performed in non-tropical regions (Menkis et al. 2014).

MOTUs classified in Archaeorhizomycetes were found in all the localities (Fig. 2B). Archaeorhizomycetes were most abundant in the wet forests El Yunque (Cubuy) and Maricao, followed by moist forest Guajataca, then dry forests Guanica and Boqueron (Figs. 2B, 3). Only a few reads were classified as Archaeorhizomycetes in samples from the littoral locality Isabela dominated by introduced *Casuarina* trees. In contrast to other Montane Wet localities, few reads were classified as Archaeorhizomycetes from the Montane wet forest site El Yunque (El Cacique). We do not have an explanation for the low numbers of Archaeorhizomycetes from the El Cacique site, though this could be due to type of soil or local variations. Current sampling could not resolve whether locality or ecosystem had a greater effect on the distribution of Archaeorhizomycetes in the island. We found that 84 MOTUs (45.7 %) differed in abundances among localities (Bonferroni *P* < 0.01; Supplementary Table S1) while 58 MOTUs (31.5 %) differed among ecosystems (Bonferroni *P* < 0.01; data not shown). The high variation in Archaeorhizomycetes abundance among localities is in accordance with earlier observations of local variation of Archaeorhizomycete (Porter et al. 2008; Rosling et al. 2013). Species abundances have been suggested to be affected by biotic and abiotic factors including type of vegetation, soil horizon, season and pH (Rosling et al. 2013).

Another abundant Ascomycota MOTU was MOTUHB 5, identified as *Phialocephala* (Vibrisseaceae). This MOTU occurred across all localities (Figs. 2B, 3; Supplementary online File S1), but was particularly abundant in littoral Isabela, which is dominated by introduced *Casuarina* trees. Species belonging to this genus are soil-and root-inhabiting fungi commonly found in alpine and boreal ecosystems in association with pine roots (Jacobs et al. 2003). High abundance of *Phialocephala* MOTUHB 5 in this locality could be explained as a result of association with mycorrhizal activity in *Casuarina* roots (Wang and Qiu 2006). Other studies have also reported *Phialocephala* species as common root-associated fungi (Bougoure and Cairney 2005) and important decomposers of organic matter in soil in tropical regions (DeAngelis et al. 2013).

Similar to the ascomycete *Phialocephala*, the abundance of Glomeromycota may reflect an association with *Casuarina* roots, with the consistency of abundance due to the local ubiquity of *Casuarina*. Whether our results reflect the actual abundance of Glomeromycota at this and other sites is unclear, given the known difficulties in amplifying ITS regions from this group of fungi in general (Hart et al. 2015).

### 4.3. Soil-inhabiting Basidiomycota in Puerto Rico

We identified basidiomycete taxa known to contain EMF, e.g., Inocybaceae, Entolomataceae, Sebacinaceae (Fig. 2C, Supplementary online File S1). EMF basidiomycete genera, *Boletus, Chantharellus, Lactarius, Suillus, Telephora, Russula* and *Scleroderma,* have previously been reported in Puerto Rico associated with the plant families Polygonaceae, Nyctaginaceae and Fabaceae (Miller et al. 2000) as well as other genera in the rhizosphere of native plants on Tenerife in the Canary Islands (Zachow et al. 2009).

The strong local variation we observed in the basidiomycete communities of Puerto Rico has also been observed in other tropical as well as boreal ecosystems (Tedersoo et al. 2012; McGuire et al. 2013). Such strong local variation could be an effect of the presence of clusters of trees associated with EMF that drastically change the local fungal community (McGuire et al. 2013).

### 4.4. Diversity of soil-inhabiting fungal community across localities and ecosystems

Based on the observed MOTU and Chao-1 richness, the moist Guajataca has the highest richness of soil fungal MOTUs and El Yunque (El Cacique) the lowest (Fig. 4). The Shannon diversity index, which takes into account abundance and evenness, suggest that protected localities (national and state forests) seem to host a higher number of soil fungal species in comparison to monoculture stands, such as Isabela, which has low plant diversity and is dominated by introduced *Casuarina* trees (Fig. 4). The low abundance and diversity of soil fungi at Isabela is also reflected by the rarefaction analysis with lower diversity Shannon index values (Figs. 4). Above we have discussed the dominance of exotic plant species *Casuarina* as a possible reason for the particular taxonomic representation of soil fungi at Isabela.

## 5. Conclusion

Soil fungal communities in Puerto Rico are organized similarly to other mature tropical and temperate soil fungal communities, with the Ascomycota dominating, followed by the Basidiomycota. In particular, we have shown that Archaeorhizomycetes is one of the most MOTU rich classes in many localities; this was possible to detect only after addressing known primer and amplification biases. Soil fungal community composition also varies significantly among localities and ecosystems, with just 26 MOTUs detected at all seven localities; all of these are common soil decomposer ascomycetes. Basidiomycetes were mostly saprophytic and they had a more local distribution compared to Ascomycetes. MOTU accumulation analysis showed that we have characterized the majority of fungal taxa present in each sample and locality. This reflects our comprehensive methodology, which includes in situ DNA extraction and the use of deep sequencing output obtained from the Ion Torrent platform.

## Disclosure

The authors declare no conflicts of interest. All the experiments undertaken in this study comply with the current laws of the country where they were performed.

## Acknowledgements

This work was funded by the Swedish Research Council VR and the Carl Trygger Foundation for Scientific Research. We thank Stefan Bertilsson, Department of Limnology, Uppsala University, for providing access to the CFR96 Touch^TM^ Real/Time PCR Detection system. We acknowledge the Departamento de Recursos Naturales y Ambientales of Puerto Rico for granting a collecting permit (2014-IC-082) to MJC.

---

**Supplementary Fig. S1:**
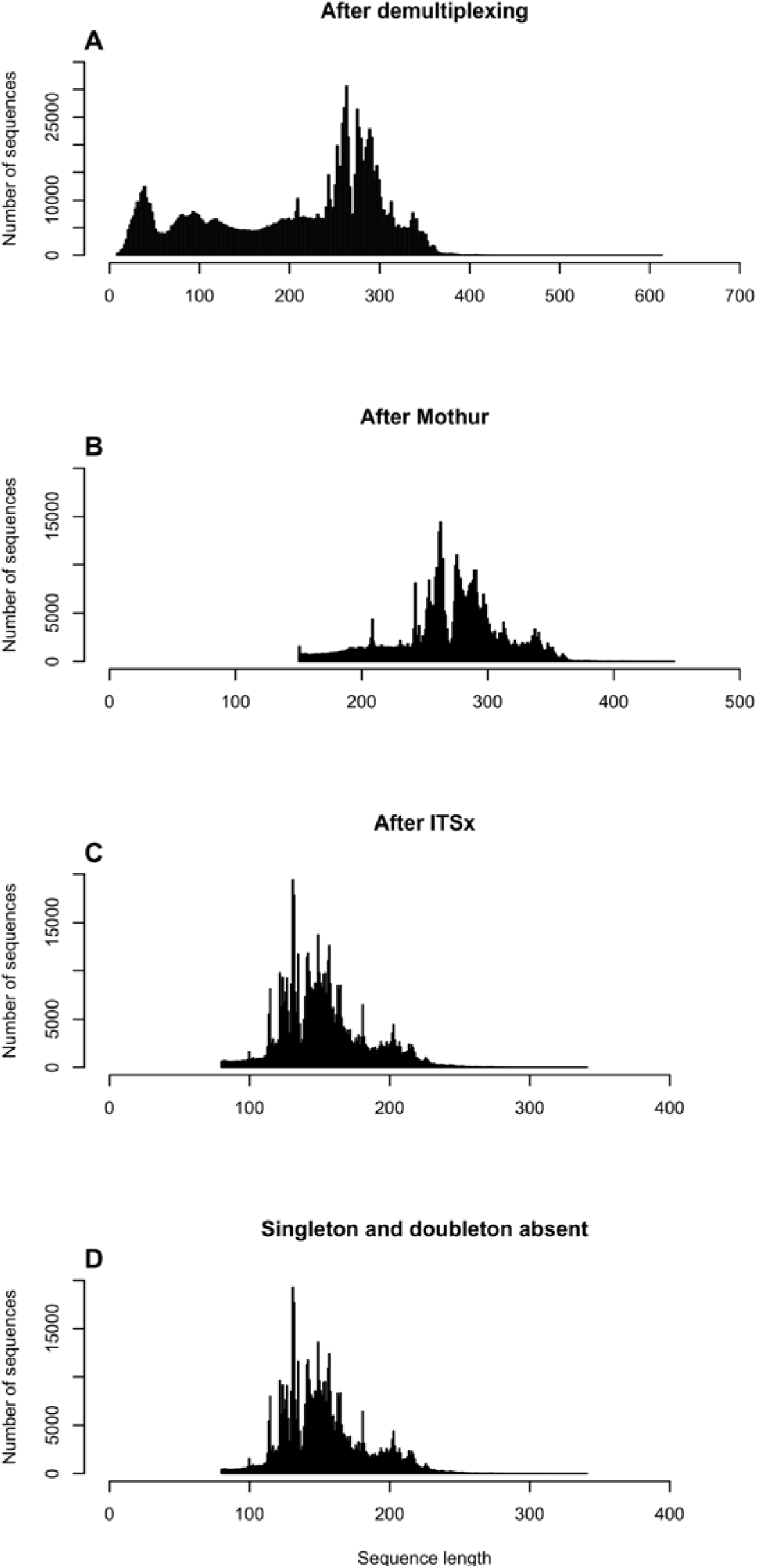
Read length frequency for all the samples: A) after demultiplexing; B) after quality filtering using mothur; C) after ITSx trimming and removal of nonfungal sequences; D) final read set after removal of singletons and doubletons.

**Supplementary Figure S2:**
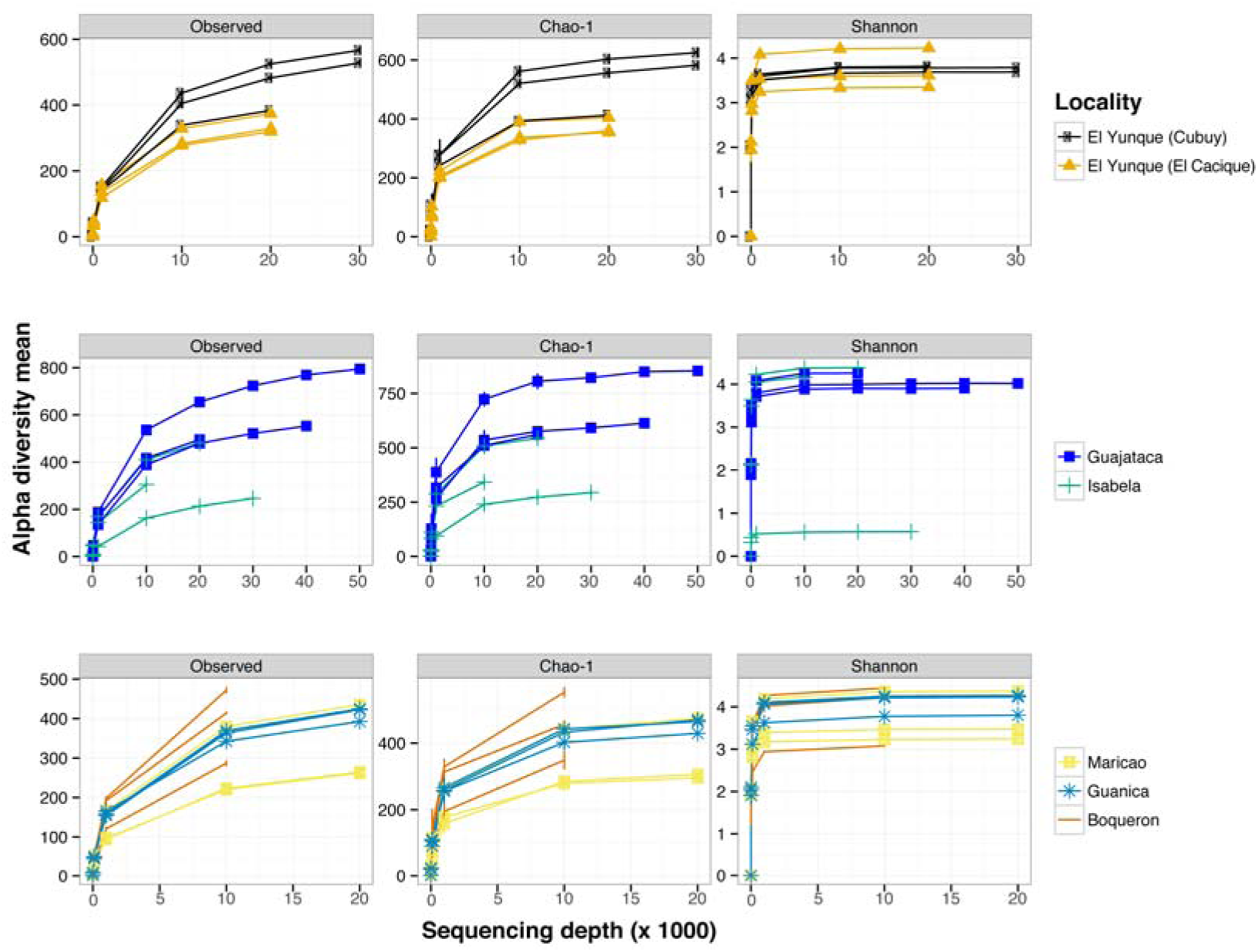
MOTU accumulation curve of alpha diversity by samples among observed species and the Chao-1 and Shannon diversity indices.

**Supplementary Figure S3:**
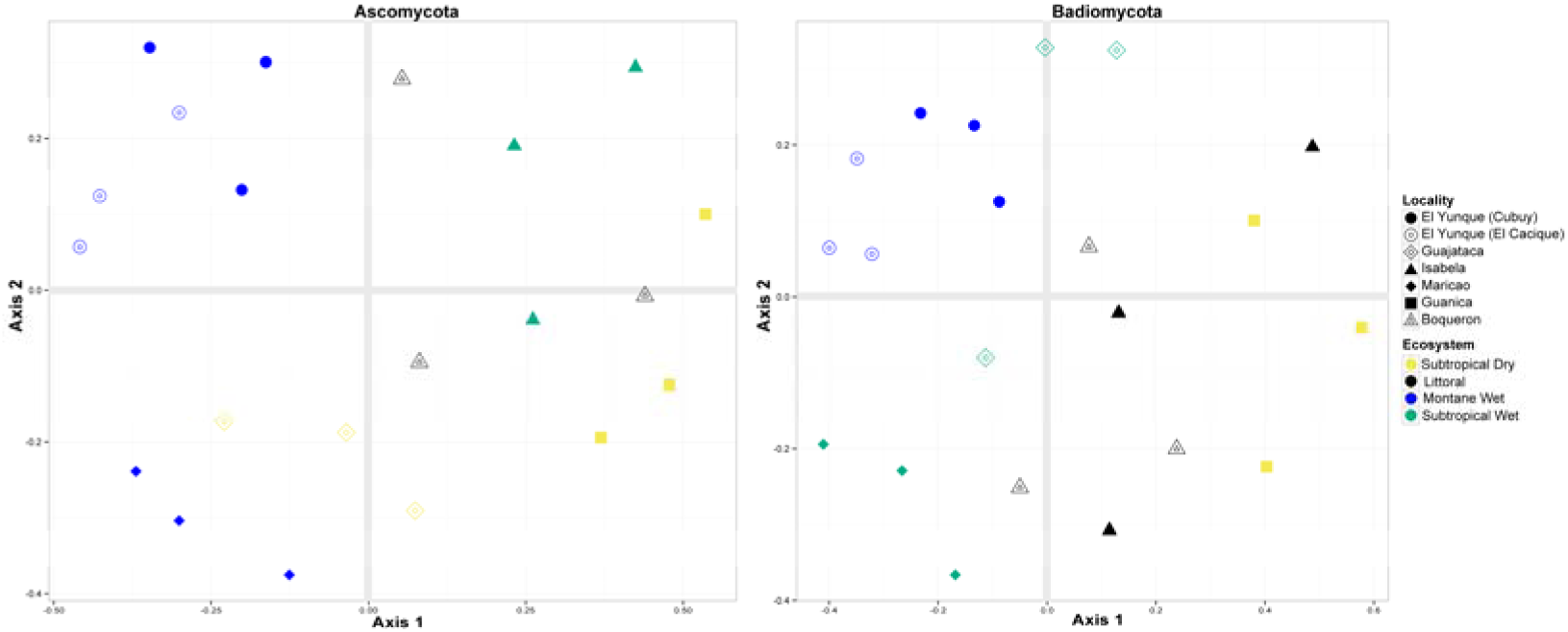
Non-metric multidimensional scaling (NMDS) of all Ascomycota MOTUs (left) and Basidiomycota MOTUs (right) in each sample, characterized by locality and ecosystem.

**Supplementary Fig. S4:**
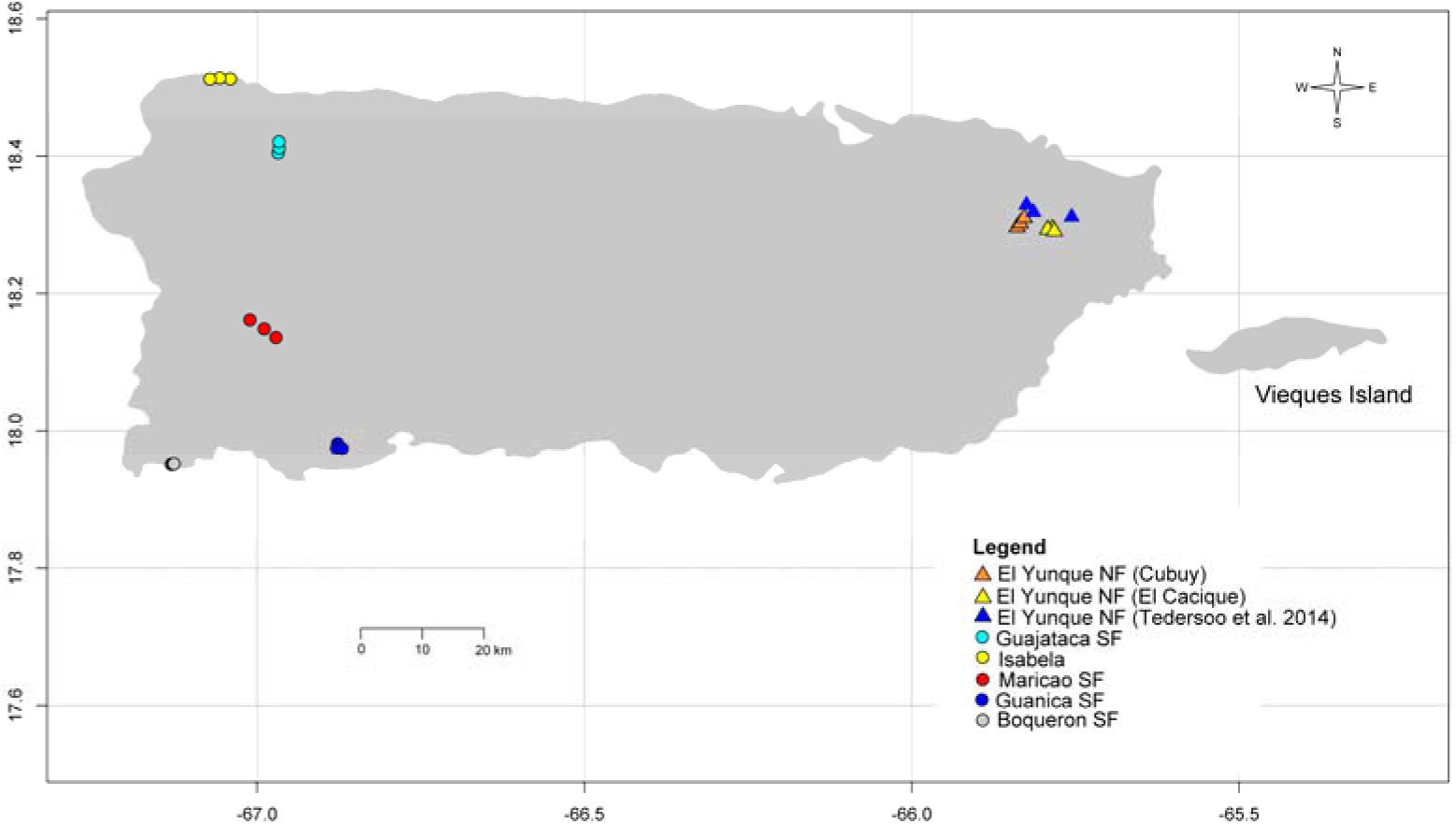
Map of Puerto Rico showing localities of sampling sites together with the El Yunque sampling sites from Tedersoo et al. (2014).

**Supplementary Table S1:**
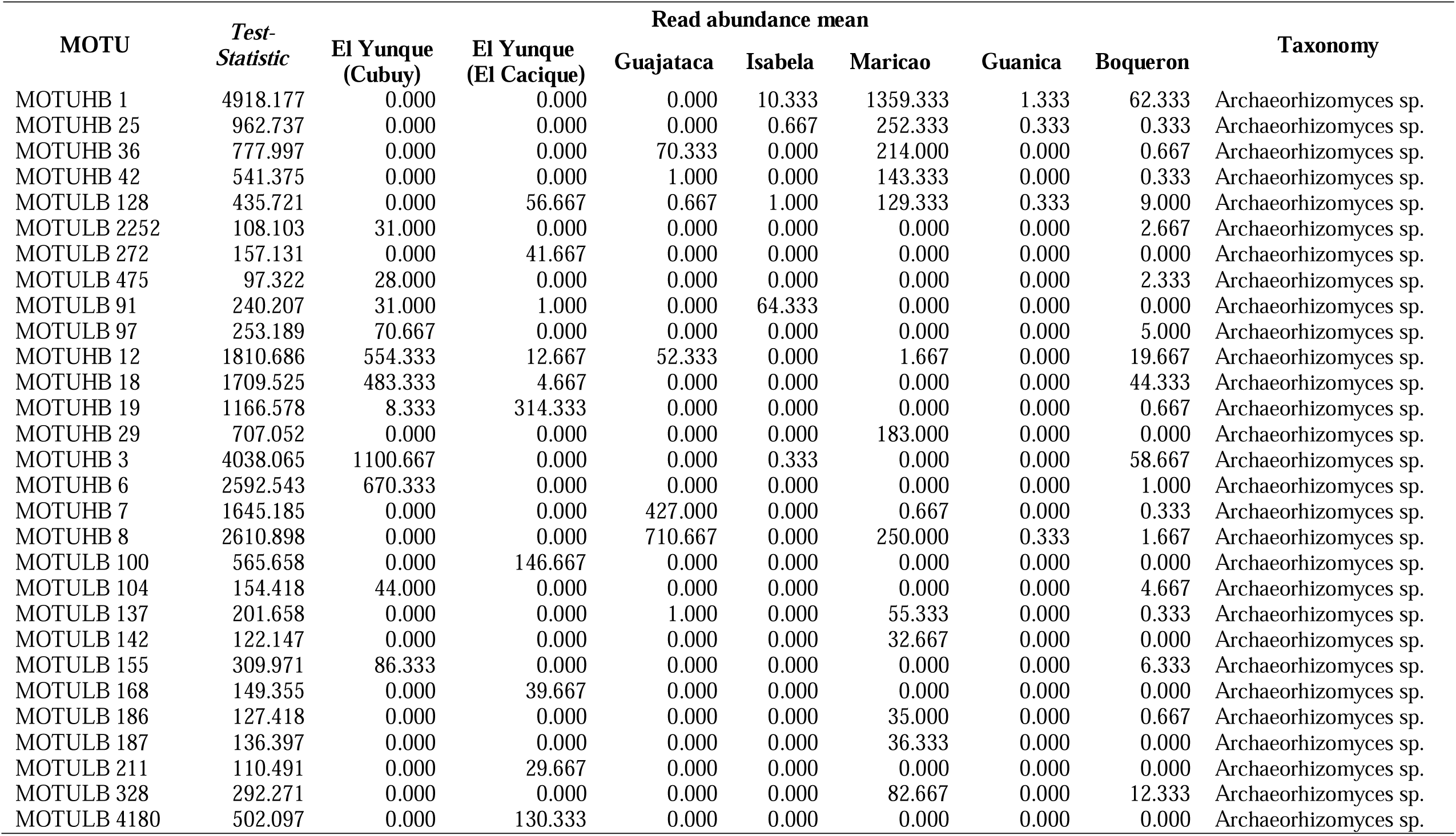
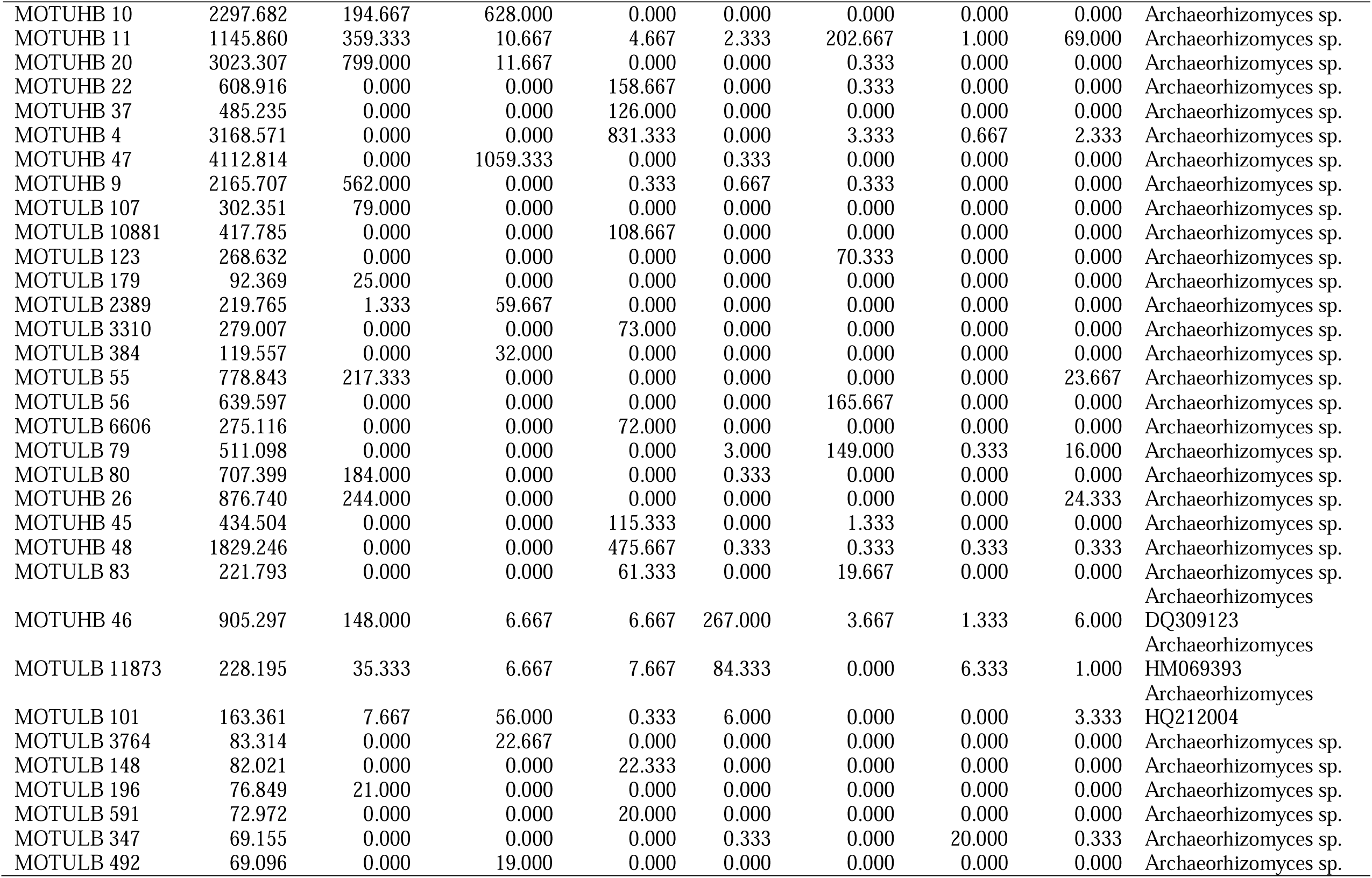
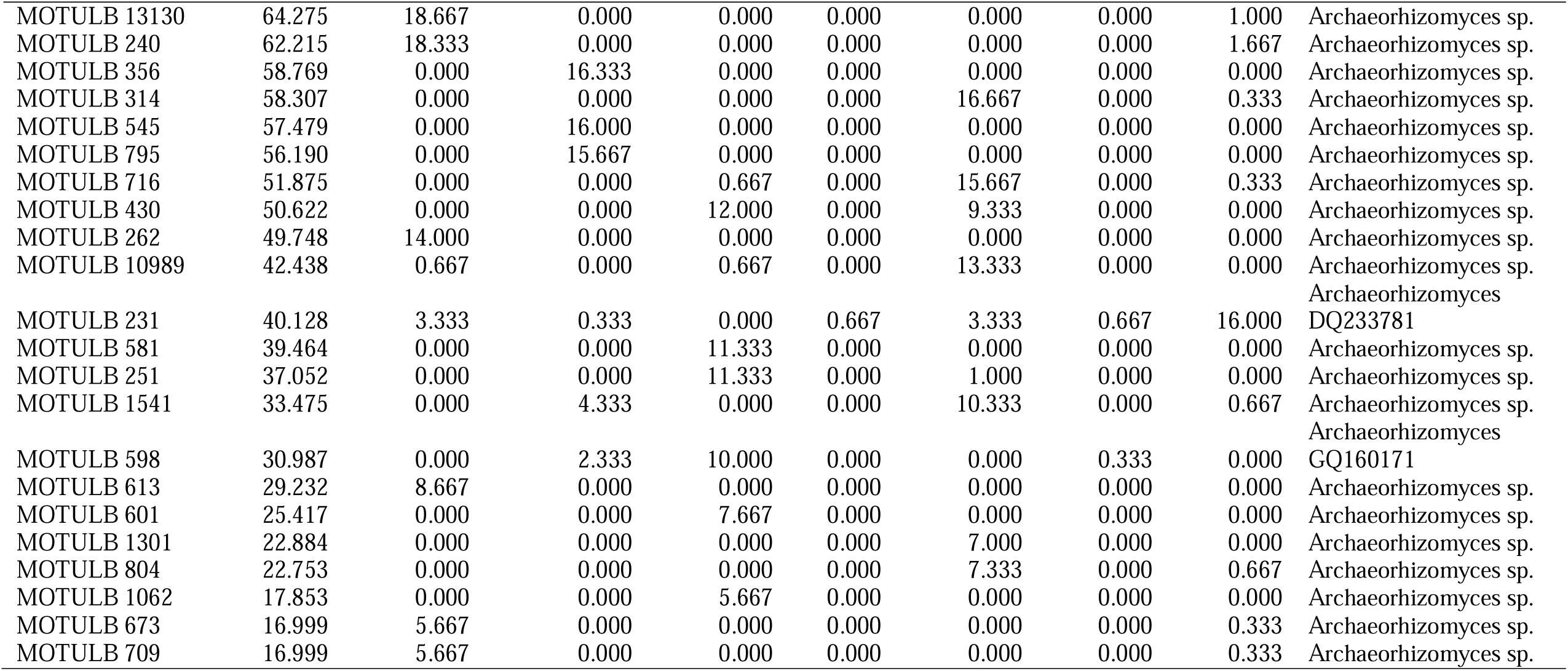
g-Test results for Archaeorhizomycetes across localities. The probabilities of the *post hoc* statistical test of False Discovery Rate (FDR) and Bonferroni correction are also included. For all MOTUs the *p* < 0.001 and FRD *p* < 0.001 and Bonferroni *p* < 0.001.

